# Divergence in female damselfly sensory structures is consistent with a species recognition function but shows no evidence of reproductive character displacement

**DOI:** 10.1101/285833

**Authors:** Alexandra A. Barnard, John P. Masly

## Abstract

Males and females exchange signals prior to mating that convey information such as sex, species identity, or individual condition. In some animals, tactile signals relayed during physical contact between males and females before and during mating appear to be important for mate choice and reproductive isolation. This is common among odonates, when a male grasps a femalexs’s thorax with his terminal appendages prior to copulation, and the female subsequently controls whether copulation occurs by bending her abdomen to complete intromission. It has been hypothesized that mechanosensory sensilla on the female thoracic plates mediate mating decisions, but is has been difficult to test this idea. Here, we use North American damselflies in the genus *Enallagma* (Odonata: Coenagrionidae) to test the hypothesis that variation in female sensilla traits is important for species recognition. *Enallagma anna* and *E. carunculatum* hybridize in nature, but experience strong reproductive isolation as a consequence of divergence in male terminal appendage morphology. We quantified several mechanosensory sensilla phenotypes on the female thorax among multiple populations of both species and compared divergence in these traits in sympatry versus allopatry. Although these species differed in features of sensilla distribution within the thoracic plates, we found no strong evidence of reproductive character displacement among the sensilla traits we measured in regions of sympatry. Our results suggest that species-specific placement of female mechanoreceptors may be sufficient for species recognition, although other female sensory phenotypes might have diverged in sympatry to reduce interspecific hybridization.

## 1 Introduction

For sexual species, maintenance of species boundaries relies on reproductive isolation (RI) between recently diverged species (Mayr, 1942). Premating reproductive isolating barriers, including behavioral or sexual isolation, often evolve earlier in the speciation process than postmating barriers in a variety of animal taxa (*e.g.*, McMillan et al., 1997; Price and Bouvier, 2002;Mendelson and Wallis, 2003; Dopman et al., 2010; Sánchez-Guillén et al., 2012; Williams and Mendelson, 2014; Castillo et al., 2015; Barnard et al., 2017). Behavioral isolation requires that mate recognition signals and/or preferences diverge between populations, which ultimately results in the ability of individuals to discriminate conspecifics from heterospecifics. Species recognition signals may rely on a variety of sensory modalities such as color (Wiernasz and Kingsolver, 1992; Sætre et al., 1997; Jiggins et al., 2001; Boughman et al., 2005; Kronforst et al., 2006; Williams and Mendelson, 2014), courtship behavior (Stratton and Uetz, 1986), sound/vibration (Ewing and Bennet-Clark, 1968; Wells and Henry, 1998; Shaw, 2000; Gerhardt and Huber, 2002; Arthur et al., 2013), and volatile chemicals (Coyne et al., 1994; Noor and Coyne, 1996; Trabalon et al., 1997; Rafferty and Boughman, 2006). Often, multiple signals act in concert to affect species recognition (*e.g.*, Costanzo and Monteiro, 2007; Girard et al., 2015).

Although much is known about the importance of visual, auditory, and chemical signals and responses in sexual communication and species recognition, we know relatively little about other sensory modalities that may have strong effects on individual mating decisions. Tactile signals have been hypothesized to contribute to mating decisions (Mendelson and Shaw, 2012), but it is unclear whether tactile cues could represent a primary species recognition signal, given that visual, auditory, and chemical cues usually act earlier during the mating sequence.

Research on the prevalence of tactile signals in mating decisions is limited (Coleman, 2008) partly because of the experimental challenge it poses: whereas other sensory modalities present male signals to a focal female from a distance, studying female preference for tactile cues requires contact between males and females, which is not always easily achieved or quantified under controlled conditions.

Despite this challenge, understanding the role of tactile signals along the continuum between intraspecific mate choice and interspecific RI is important because it broadens our understanding of the causes and consequences of a common pattern in nature — the rapid divergence of male genital morphology between species. It has been suggested that rapid genital differentiation can cause RI (Dufour 1844), although mechanical incompatibilities between heterospecific male and female genitalia do not appear to be a common cause of RI (Shapiro and Porter, 1989; Masly, 2012; Simmons, 2014). However, observations both within (Eberhard, 1994; Edvardsson and Göran, 2000; Briceño and Eberhard, 2009a; Briceño and Eberhard, 2009b; Frazee and Masly, 2015) and between species (Patterson and Thaeler Jr, 1982; Robertson and Paterson, 1982; Eberhard, 1992; Coyne, 1993; Barnard et al., 2017) suggest that male reproductive structures may convey tactile information to females that affects their subsequent behavior and/or physiology. Although female genital structures often appear invariant among closely related species (Shapiro and Porter, 1989), subtle morphological differences (*e.g.*, Kamimura and Mitsumoto, 2011; Yassin and Orgogozo, 2013) could enable females to detect variation among males’ morphology. Female variation in detection ability could also occur in signal processing at the level of neurons, neural networks, and/or in the distribution and morphology of sensory structures that receive male tactile signals. These sensory structures may exist not just in the female genitalia or reproductive tract, but in any region of the female that receives contact from male structures.

Female sensory structures that reside in body regions that contact species-specific male structures during mating have been documented in several arthropods, including flies (Eberhard, 2001; Ingram et al., 2008) and damselflies (Córdoba-Aguilar, 1999, 2002, 2005; Robertson and Paterson, 1982). Other studies have demonstrated that tactile cues from male grasping organs influence female mating responses, either via experimental manipulation of male structures and desensitization of females (Eberhard, 2002; Briceño et al., 2007; Briceño and Eberhard, 2009a; Eberhard, 2010; Myers et al., 2016), or via comparison of female behavior when females are grasped by males with varying terminal appendage morphologies (Sánchez-Guillén et al., 2012; Sánchez-Guillén et al., 2014; Barnard et al., 2017). Premating tactile isolation may also be important in vision-limited vertebrates. For example, contact cues via the lateral line system may influence female mate choice in a cavefish (Plath et al., 2004; but see Rüschenbaum and Schlupp, 2013).

Tactile signals appear to be a significant cause of RI in Zygoptera, the damselfly suborder of Odonata (Krieger and Krieger-Loibl, 1958; Loibl, 1958; Robertson and Paterson, 1982; Corbet, 1999). Concentrations of cuticular mechanoreceptors (sensilla) on the female thorax have been described in several coenagrionid damselfly genera. The morphology of these sensilla is consistent with a mechanosensory function and does not indicate that they conduct signals related to olfaction, hygroreception, or temperature reception (McIver, 1975; Robertson and Patterson, 1982). These sensilla reside in areas where males’ grasping appendages contact the female thorax before and during mating, which has led to speculation that they allow females to evaluate male morphologies and discriminate conspecific from heterospecific males (Jurzitza, 1974, 1975; Tennessen, 1975; Robertson and Paterson, 1982; Battin, 1993a, 1993b). Each mechanoreceptor is associated with a single sensory neuron (McIver, 1975; Kiel, 1997). The thoracic sensilla thus represent a spatial matrix that can transmit signals to the female central nervous system based on the pattern in which the sensilla are stimulated. Greater numbers of these receptors are expected to enhance a female’s sensory resolution by increasing the combinatorial complexity of tactile signals that she can perceive. For example, if a female possesses 25 sensilla, and each sensillum has two response states (“on” if contacted and “off” if not contacted), then the number of unique tactile patterns that the female could distinguish is 2^25^ = 3.4 × 10^7^. A female that possesses just one additional sensillum would be able to distinguish among roughly twice as many tactile patterns (2^26^ = 6.7 × 10^7^). Should individual sensilla respond to quantitative variation in touch (rather than a binary response), this would dramatically increase the number of response states and therefore further enhance tactile acuity (*e.g.*, Gaffin and Brayfield, 2017). Female damselfly thoracic sensilla thus present an external, quantifiable phenotype in which to investigate the mechanistic basis of tactile stimuli and female mating decisions.

The North American damselfly genus *Enallagma* includes several recently diverged species that often co-occur in the same habitats (Johnson and Crowley, 1980; McPeek, 1998), and do not engage in premating courtship (Fincke et al., 2007; Barnard et al., 2017) or use chemical cues for mate selection (Rebora et al., 2018). A female’s first opportunity to assess a potential mate occurs when the male uses his terminal appendages to grasp the mesostigmal plates on the female thorax to form “tandem”, the premating position. The male superior grasping appendages (cerci) have species-specific morphologies, and differences in the morphology of these structures are the primary cause of RI in this genus (Paulson, 1974; Barnard et al., 2017). Two species, *E. anna* and *E. carunculatum*, occasionally hybridize in nature to produce males and females with reproductive structure morphologies that are intermediate to each of the pure species (Miller and Ivie, 1993; Donnelly, 2008; Johnson, 2009; Barnard et al., 2017). Females of both pure species discriminate strongly against both heterospecific and interspecific hybrid males that take them in tandem, which shows that female *E. anna* and *E. carunculatum* can detect not only large differences in male cercus morphologies, but also more subtle differences such as those that distinguish conspecific and hybrid males (Barnard et al., 2017).

Because it appears that mesostigmal sensilla mediate species recognition, they might be expected to show signs of reproductive character displacement (RCD): increased divergence of traits involved in RI in regions of sympatry between *E. anna* and *E. carunculatum* relative to regions of allopatry (Brown and Wilson, 1956; Howard, 1993; Pfennig and Pfennig, 2009). RCD can manifest phenotypically as divergence in either signaling traits or mate preferences, in which sympatric females display stronger discrimination against heterospecific males than do allopatric females of the same species (*e.g*., Gerhardt, 1994; Gabor and Ryam, 2001; Albert and Schluter, 2004; Wheatcroft and Qvarnström, 2017). This strengthening of preference in sympatry may evolve via direct selection on adult prezygotic phenotypes, or via reinforcement, where selection against interspecific hybrids gives rise to selection for enhanced premating isolation between species (Dobzhansky, 1937). *Enallagma anna* and *E. carunculatum* can interbreed, but their hybrids experience significantly reduced fitness (Barnard et al., 2017). Female *Enallagma* experience frequent mating attempts from heterospecific males where both species co-occur (Paulson, 1974; Fincke et al., 2007; Barnard et al., 2017). These findings suggest that in sympatry, females may experience selection for stronger species discrimination ability. Studies of several *Enallagma* species (not including *E. anna* or *E. carunculatum)* have revealed that male cercus shape varies little among populations, even across large geographical regions (McPeek et al., 2011; Siepielski et al., 2018). *Enallagma anna* and *E. carunculatum* appear to show similar patterns, at least in the western part of their distributions (A. Barnard, unpublished data). It is possible, however, that females in sympatry with other species are more sensitive to variation among males than are females of the same species in regions of allopatry, and this variation in sensitivity may be reflected in female sensilla traits.

Here, we use sensilla number, density, and location as proxies for female preference, to test the hypothesis that variation in female sensilla phenotypes supports a function in species recognition. We tested this hypothesis by quantifying sensilla on the mesostigmal plates of a large set of *E. anna* and *E. carunculatum* females from multiple populations across the western United States and comparing phenotypes of each pure species from sympatric and allopatric populations to identify patterns consistent with RCD. We predicted that in sympatric populations, females would possess higher sensilla numbers, higher sensilla density, and/or different spatial distributions of sensilla within their mesostigmal plates when compared to females from allopatric populations.

## Materials and methods

### Population sampling

We measured the sensilla traits of 29 *E. anna* females across 13 populations, and 74 *E. carunculatum* females across 19 populations (Fig. 1, Table 1). We classified each population as allopatric, locally allopatric, or sympatric. Sympatric populations are those where *E. anna* and *E. carunculatum* co-occur temporally as well as spatially. Because *E. anna’s* geographic range falls completely within *E. carunculatum*’s range, only *E. carunculatum* has completely allopatric populations. We designated populations as locally allopatric at sites within the area of range overlap, but where only one species is known to occur based on occurrence data from OdonataCentral (Abbott, 2016). Although some specimens were collected as early as 1945, the majority of samples (82 of 103) we studied were collected between 2012 and 2016.

**Figure 1.**
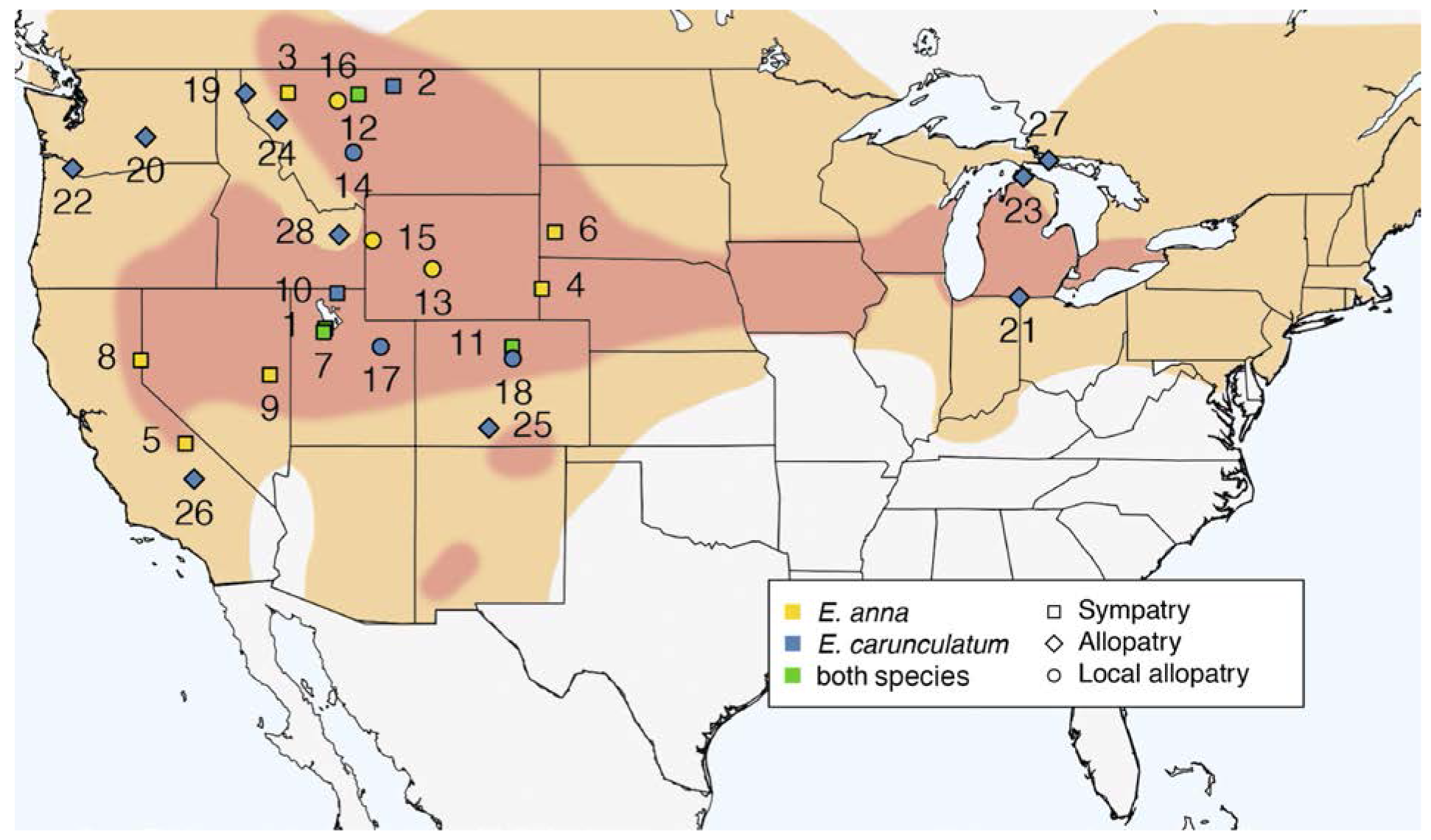
Sampling sites and species ranges. *Enallagma anna’s* geographic range (red) occurs within *E. carunculatum*’s geographic range (orange). Names of sites associated with each number are described in Table 1. Symbol color indicates the species sampled and symbol shape indicates the population type. (Species ranges are adapted from Johnson, 2009; Paulson, 2009, 2011).

**Table 1.**
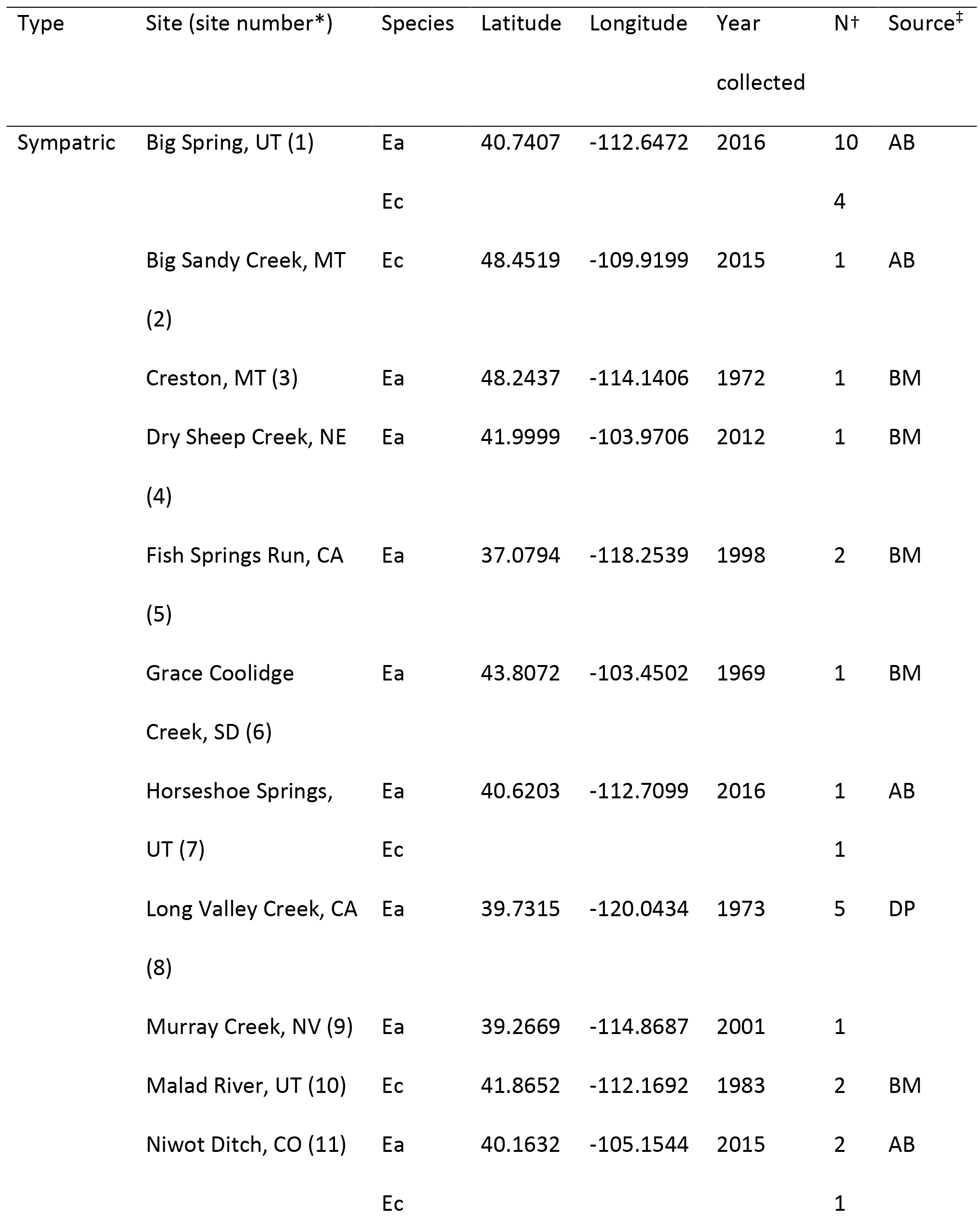
Sampling sites for *E. anna* and *E. carunculatum* populations.

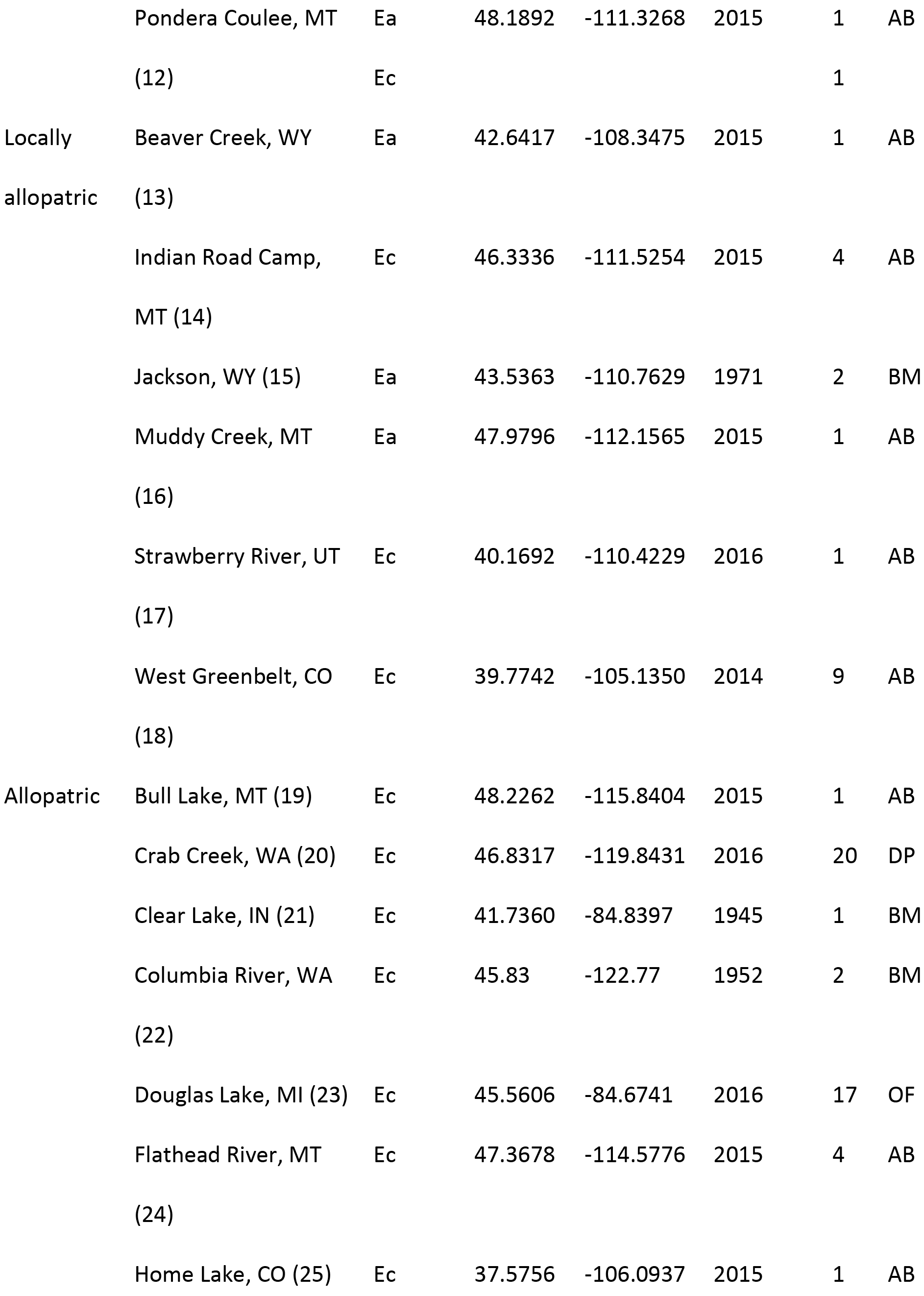

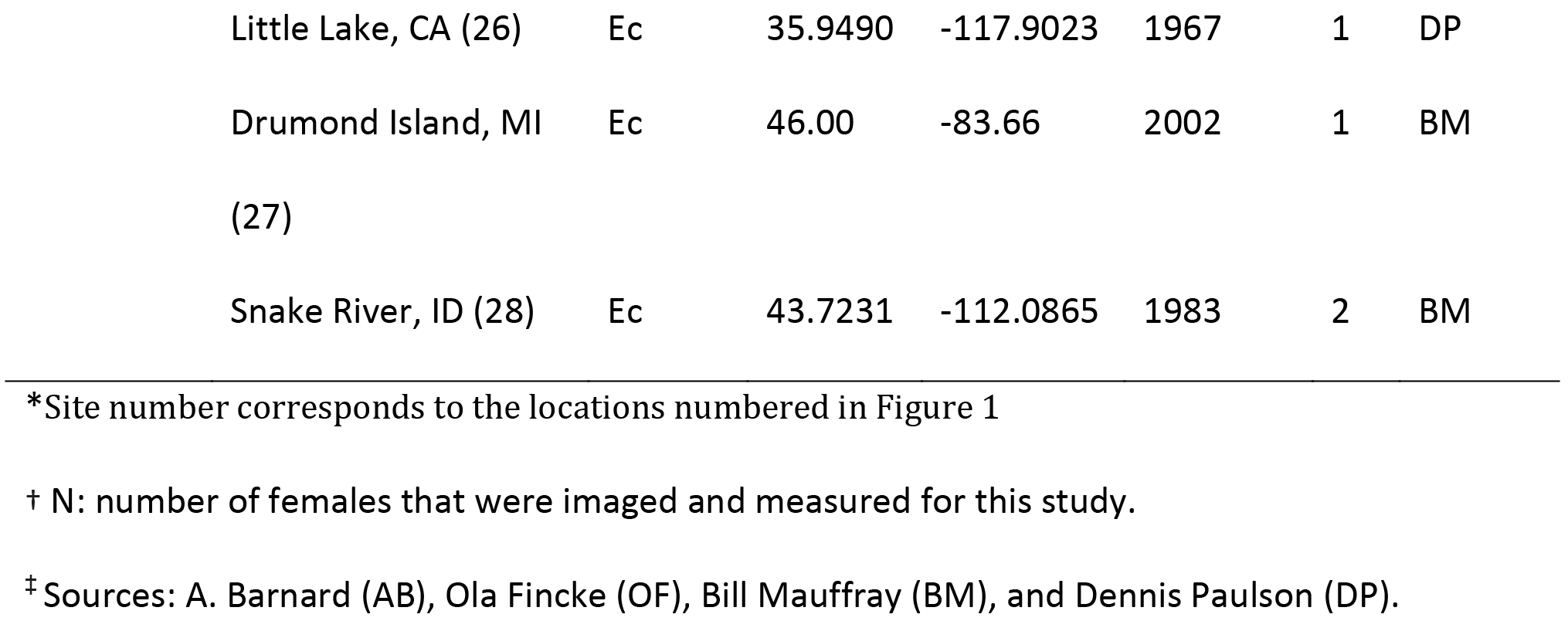

### Trait imaging and quantification

We photographed each damselfly using a Nikon D5100 camera (16.2 MP; Nikon Corporation, Tokyo, Japan). We dissected the ventral thoracic cuticle from each female using forceps and imaged the mesostigmal plates using scanning electron microscopy. Specimens were mounted on aluminum stubs with carbon tape, sputter-coated with gold-palladium, and imaged at, 200X magnification and 3kV using a Zeiss NEON scanning electron microscope.

To avoid any potential bias during measurements, we blind-coded image files before measuring all traits. We measured abdomen length (abdominal segments 1-10, excluding terminal appendages) on the full-body photos as an estimate for body size using the segmented line tool in ImageJ (Abramoff et al., 2004). We quantified sensilla traits on the right mesostigmal plate of each female damselfly unless the right plate was dirty or damaged, in which case we quantified the left plate (n = 20). Sensilla counts on a subset of 57 females showed that left plate and right plate sensilla counts were highly correlated within individual females (r = 0.85). In cases where we quantified the left plate, we flipped the image horizontally, so it was in the same orientation as a right plate. We standardized the position of the mesostigmal plate in each image by cropping and rotating the image so that the lower medial corner of the plate was in line with the lower left corner of each image. We counted sensilla and obtained their *x* and *y* coordinates in ImageJ using the multi-point selection tool. We traced an outline around the plate image, excluding the lateral carina (Fig. S1), using a Wacom Cintiq 12WX tablet and stylus (Wacom, Saitama, Japan) and the freehand selection tool in ImageJ. This procedure produced *x* and *y* coordinates that described the plate outline. We performed all measurements twice for each specimen. Measurements across the two technical replicates were highly correlated (*r*_abdomen_ = 0.95, n = 78; *r*_count_ = 0.95, n = 103; *r*_plate area_ = 0.99, n = 86), so we used the mean trait values of the two replicates in subsequent analyses. Seventeen samples were imaged at angles that allowed counting of the sensilla, but distorted the plate shape or distances between the sensilla. Those samples are included in analyses of sensilla number, but were not included in the analyses of sensilla density or distribution.

### Sensilla trait analyses

We conducted all morphometric and statistical analyses using R v. 3.4.1 (R Core Team, 2015). We used the mesostigmal plate outline coordinates to calculate each plate’s two dimensional area. To calculate the area of the sensilla-covered region of each plate, we generated a polygon connecting the coordinates of the outermost sensilla and calculated the area within this outline. We determined the proportion of each plate that was covered by sensilla by dividing the sensilla area by total plate area. We calculated sensilla density in two ways. First, we divided sensilla number by the area of the sensilla-covered region. This measures the number of sensilla that occur in a particular area, but does not capture the relative arrangement of sensilla within that area. Second, we computed the nearest neighbor distances among all sensilla within each plate based on their *x* and *y* coordinates and then calculated the mean and median nearest neighbor distances between the sensilla for each female. Nearest neighbor mean and median distances were highly correlated (*r*_*E. carunculatum*_ = 0.83; *r*_*E. anna*_ = 0.81), so we used the mean values for these measures in our analyses.

To determine whether larger females possess more sensilla, we regressed sensilla number against abdomen length. We found no significant relationship between these traits in either species (*E. anna:* R^2^_adj_ = -0.007, F_1,25_ = 0.82, *P* = 0.737; *E. carunculatum: R*^2^_adj_ = 0.01, *F*_1,48_ = 0.52, *P* = 0.47). We thus present the results that compare sensilla counts without correcting for differences in body size.

### Sensilla spatial analyses

To quantify sensilla distributions within each plate, we generated kernel density estimates (KDEs) for populations with at least four sampled individuals (two *E. anna* and six *E. carunculatum* populations) using the R package ks (Duong, 2016). First, we randomly selected one of the two replicate sets of sensilla and plate outline coordinates for each female. To prepare the coordinate data for KDE analyses, we concatenated the sensilla and plate coordinates for each female and adjusted all plate outlines to have an area of one. This standardized each set of sensilla coordinates for size, while maintaining their relative positions within each plate. Next, we translated each set of coordinates to place the origin of the coordinate system at the plate outline’s centroid. We concatenated sensilla coordinates for all females sampled within each population to compute a representative KDE for each population.

Within each species, we conducted pairwise tests to compare each population’s KDE against every other population using the function kde.test with the default settings. This test returns a *P*-value that reflects the probability of generating the two respective KDEs from the same distribution of points. Because we performed multiple pairwise tests among *E. carunculatum* populations, we adjusted the resulting P-values using the false discovery rate (Benjamini & Hochberg, 1995).

We generated an average plate outline for each population on which to visualize the KDEs. The total number of coordinates that describe each plate outline varied among females, ranging from 647-1078 for *E. anna* and 688-1028 for *E. carunculatum.* We standardized the number of coordinates representing each plate by retaining the points for each of the upper and lower medial corners and randomly sampled 198 points in between. We then treated each of these, 200 points as landmarks (the corners represented fixed landmarks and the remaining points were designated as sliding semilandmarks) and used the R package geomorph (Adams and Otarola-Castillo, 2013) to perform general Procrustes analysis (Rohlf, 1999) and obtain an average two-dimensional plate shape for each population.

### Statistical analyses

Some populations were well-sampled whereas others were represented by a single female (Table 1). To avoid psedoreplication, for each population with N > 1, our analyses of sensilla number, density, and area of each mesostigmal plate covered by sensilla used the population mean of each trait value, so that each population was represented by a single measurement. We arcsin transformed proportion data prior to analysis. To compare traits between *E. anna* and *E. carunculatum*, we used Welch’s *t*-tests. We compared traits among sympatric, locally allopatric, and fully allopatric *E. carunculatum* populations using Kruskal-Wallis tests, and between sympatric and locally allopatric *E. anna* populations using Welch’s *t*-tests. To understand the relationships between sensilla number, sensilla density, and the area of the plate occupied by sensilla, we performed linear regressions between each pair of traits. Due to the limitation of single samples from some populations, we analyzed *E. carunculatum* populations in two ways: we first included all samples, then conducted a separate analysis that excluded populations with N < 4. Both analyses yield similar results; we report the results for the analysis using all samples in the main text and provide results from the subset of samples with N ≥ 4 in the supplemental material (Table S1).

## Results

### *Enallagma anna* and *E. carunculatum* females possess distinct sensilla traits

*Enallagma anna* females possessed significantly more sensilla per plate (x̄ = 49 + 2) than *E. carunculatum* females (x̄ = 28 ± 1, *t*_35.1_ = 11.13, *P* = 4.6 × 10^-13^; Fig. 2A). *Enallagma anna* females also possessed sensilla distributed over a larger proportion of each plate (*t*_39.7_ = 11.1, *P =* 8.6 x 10^-14^; Fig. 2B), and larger mean distances between sensilla (*t*_54_ = 6.7, *P* = 1.3 × 10^-8^; Fig. 2C). This ultimately results in a lower density of sensilla per unit area in *E. anna* compared to *E. carunculatum* (*t*_99.6_ = −12.96, *P* = 2.2 × 10^-16^; Fig. 2D). The sensilla also occurred in different locations on the mesostigmal plates of each species: they were more medially located in *E. anna* and more laterally located in *E. carunculatum* (Figs. 3, 4).

Both species showed a strong positive relationship between sensilla number and the absolute area of the plate occupied by sensilla (*E. anna: R*^2^_adj_ = 0.33, *F*_1,27_ = 14.71, *P* = 0.0007; *E. carunculatum: R*^2^_adj_ = 0.33, *F* _1,72_ = 37.68, *P* = 4.1 x 10^-8^). Consistent with this result, linear regressions also revealed that females with more sensilla also had a larger proportion of the plate occupied by sensilla (*E. anna: R*^2^_adj_ = 0.26, *F*_1,27_ = 10.65, *P* = 0.003; *E. carunculatum: R*^2^_adj_ = 0.20, *F*_1,65_ = 18.93, *P* = 4.4 × 10^-5^). Females with more sensilla had smaller mean distances between neighboring sensilla (*E. anna: R*^2^_adj_ = 0.11, *F*_1,27_ = 4.34, *P* = 0.046; *E. carunculatum: R*^2^_adj_ = 0.09, *F*_1,72_ = 3.80, *P* = 0.01). Overall, these results show that a greater number of sensilla was more strongly associated with a sensilla distribution that covers a larger area of the mesostigmal plate rather than a greater concentration sensilla within in a smaller area.

### *E. carunculatum* sensilla traits do not show a strong pattern of reproductive character displacement

We made several non-mutually exclusive predictions expected under RCD for the sensilla traits we measured in sympatric populations relative to allopatric populations. In particular, we predicted to observe at least one of the following phenotypic differences in sympatric females relative to allopatric females: (1) more numerous sensilla, (2) denser sensilla, (3) sensilla concentrated in different regions of the mesostigmal plates. We did not find significant differences in any of these traits between sympatric and locally allopatric *E. anna* females (Table 2). However, because our *E. anna* samples included only four females from three locally allopatric populations, we could not perform a robust comparison of *E. anna* sensilla traits between populations that do, or do not encounter *E. carunculatum.* We thus focus our analysis on comparisons between sympatric and allopatric *E. carunculatum* populations, for which we had larger sample sizes.

**Table 2.** Statistical comparison of sensilla traits in locally allopatric and sympatric *E. anna* populations.

| Trait | Mean ± s.e.m. | | *t* | d.f. | *P* |
| --- | --- | --- | --- | --- | --- |
|  | Local allopatry | Sympatry |  |  |  |
|  | (N* = 3) | (N = 10) |  |  |  |
| Sensilla number | 39.8 ± 3.8 | 48.5 ± 2.3 | -1.93 | 3.6 | 0.13 |
| Proportion plate containing sensilla | 0.67 ± 0.27 | 0.69 ± 0.15 | -0.25 | 2.8 | 0.82 |
| Sensilla density (sensilla mm<sup>-1</sup>) | 1.5 ± 0.2 | 1.6 ± 0.1 | -0.43 | 2.9 | 0.70 |
| Mean distance (μm) between sensilla pairs | , 20.0 ± 1.0 | , 19.3 ± 0.5 | 0.62 | 3.13 | 0.58 |

\* N: number of populations analyzed.

Sympatric, locally allopatric, and fully allopatric *E. carunculatum* populations did not differ significantly from one another in sensilla number (Kruskal-Wallis χ^2^_2_ = 0.69, *P* = 0.71), proportion of the mesostigmal plate covered by sensilla (Kruskal-Wallis χ^2^_2_ = 2.16, *P* = 0.34), or sensilla density (overall density: Kruskal-Wallis χ^2^_2_ = 0.12, *P* = 0.94; mean distance between sensilla: Kruskal-Wallis χ^2^_2_ = 3.53, *P* = 0.17). In addition to divergence of mean trait values, RCD can also result in reduced trait variance in sympatry without affecting the mean (Pfennig and Pfennig 2009). Sympatric *E. carunculatum* populations displayed less interpopulation variance than allopatric populations in both mean sensilla number (Figure 2A) and mean proportion of the plate covered by sensilla (Figure 2B). However, these trends were not statistically significant (sensilla number: Bartlett’s K^2^_1_ = 0.83, *P* = 0.36; proportion of plate covered by sensilla: Bartlett’s K^2^_1_ = 1.86 *P* = 0.17).

**Figure 2.**
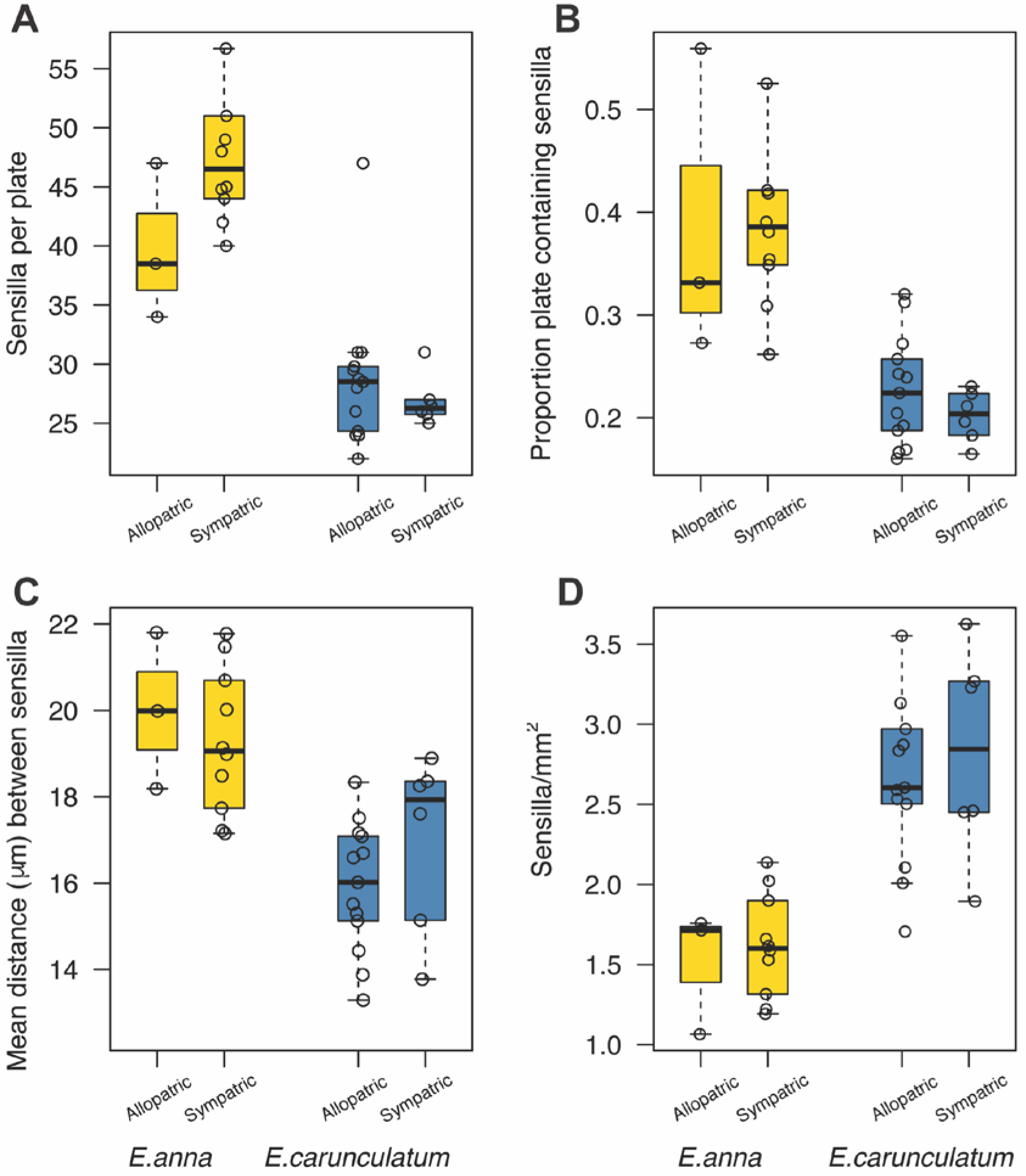
*Enallagma anna* and *E. carunculatum* sensilla traits by population type. **(A)** The number of sensilla on one mesostigmal plate. **(B)** Proportion of the plate that contains sensilla. **(C)** Mean nearest neighbor distances between sensilla. **(D)** Sensilla density in the region of the plate that contains sensilla. Within each panel, each open circle represents the mean of one population. Boxplots show the interquartile range. The line within the box shows the median and whiskers extend to the most extreme observation within 1.5 times the interquartile range.

Interestingly, although mean trait values did not differ significantly between sympatric and allopatric populations, sensilla traits displayed considerable variation within the populations we sampled. For example, within a single population, a particular female might have twice as many sensilla than another female (Fig. 3). This pattern was also observed in the *E. anna* populations we studied.

**Figure 3.**
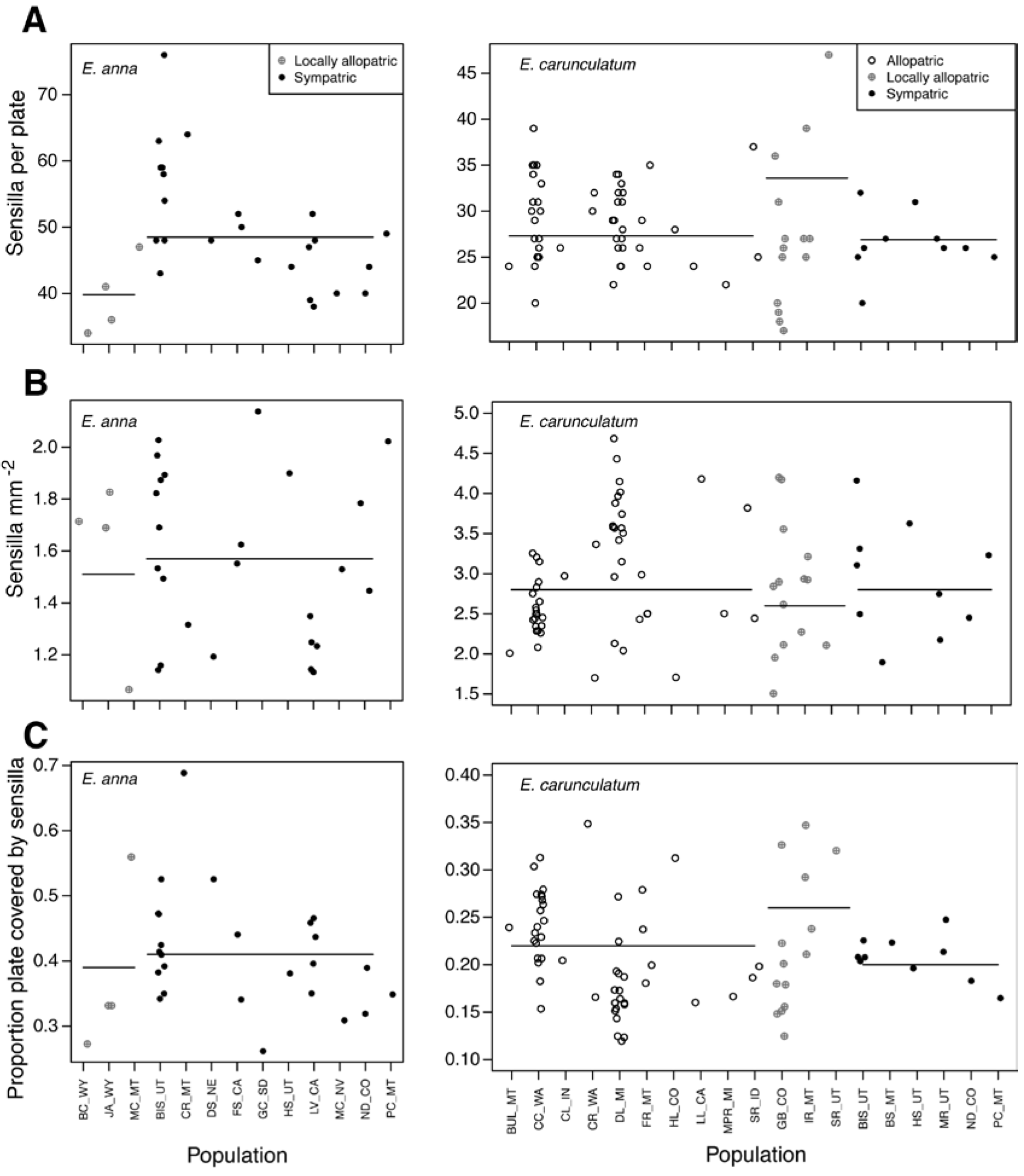
Individual trait values for sensilla number, sensilla density, and proportion of plate containing sensilla. Each symbol represents a single female, separated by population along the y-axis. Horizontal lines indicate the mean value for each population type (completely allopatric, locally allopatric, or sympatric), calculated from population means. Populations are described in Table 1.

KDE comparisons did not reveal significant differences in sensilla distributions between sympatric and allopatric *E. carunculatum* populations (Table 3). However, the analysis revealed significant differences in sensilla distributions between several pairs of allopatric *E. carunculatum* populations (Fig. 4E), which indicates that populations isolated from *E. anna* vary more among themselves than do populations sympatric with *E. anna*, which share similar sensilla patterns. This result is consistent with those described above that indicated higher variance in sensilla traits among allopatric populations compared to sympatric populations.

**Figure 4.**
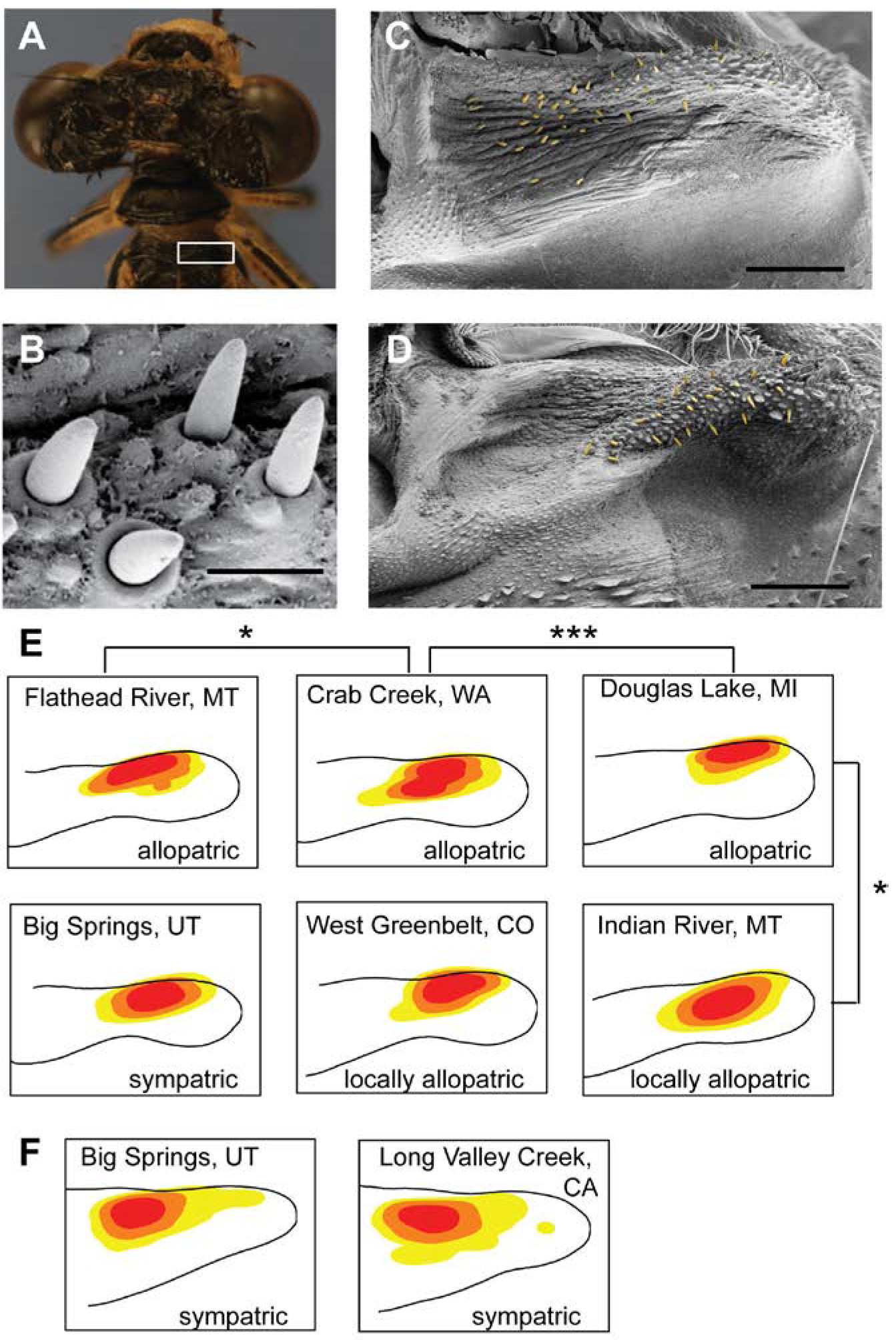
Sensilla locations. **(A)** White box indicates the location of right mesostigmal plate on the thorax. **(B)** Ultrastructural details of individual sensilla. Scale bar represents 10 μm. **(C, D)** Scanning electron micrographs show the locations of sensilla (yellow) on the mesostigmal plates of *E. anna* (C) and *E. carunculatum* (D). Scale bars represent 100 **(E, F)** Population kernel density estimates for *E. carunculatum* (E) and *E. anna* **(F)** sensilla. The shading indicates different regions of sensilla density: red represents the 75-99^th^ percentile of sensilla density, orange represents the 50-74^th^ percentile, and yellow represents the 25^th^−49^th^ percentile. Each outline represents the average mesostigmal plate shape for the population. Asterisks indicate *E. carunculatum* populations whose KDEs are significantly different (* *P* < 0.05, *** *P* < 0.001).

**Table 3.** Results of pairwise comparisons of sensilla kernel density estimates for *E. carunculatum* populations. False discovery rate-adjusted P-values are reported^t^.

|  | Big | Crab | Douglas | Flathead | West |  |  |
| --- | --- | --- | --- | --- | --- | --- | --- |
|  | Springs, | Creek, | Lake, MI | River, | Green- |  | Population |
| Population | UT | WA |  | MT | belt, CO | N* | type |
| Big Springs, UT | 1 |  |  |  |  | 4 | sympatric |
| Crab Creek, WA |  | 1 |  |  |  | 20 | allopatric |
| Douglas Lake, MI | 0.263 | **2.53e<sup>-10</sup>** | 1 |  |  | 17 | allopatric |
| Flathead River, MT | 1 | **0.0103** | 0.263 | 1 |  | 4 | allopatric |
|  |  |  |  |  |  |  | Locally |
| West Greenbelt, CO | 1 | 0.0625 | 0.3835 | 0.502 | 1 | 4 | allopatric |
|  |  |  |  |  |  |  | Locally |
| Indian River, MT | 1 | 1 | **0.0103** | 0.0625 | 0.3115 | 4 | allopatric |

\* N: number of females whose sensilla coordinates were used to calculate KDEs.
† Bold values indicate *P* < 0.05.

## Discussion

*Enallagma anna* and *E. carunculatum* females possess different numbers of sensilla in species-specific distributions on their mesostigmal plates. This result supports the idea that receptors that receive male stimuli will occur in patterns that correspond to the male organs during contact (Eberhard, 2010). An association between male morphology and female sensilla has been described for African *Enallagma* species (Robertson and Paterson, 1982), and our results show a similar pattern for two North American species. *Enallagma anna* male cerci are considerably larger than *E. carunculatum* cerci, and the observation that *E. anna* females had a larger number of sensilla compared to *E. carunculatum* females is consistent with the likelihood that *E. anna* male cerci make greater spatial contact with the mesostigmal plates.

When species make secondary contact after initial divergence in allopatry, the possible outcomes are increased species divergence (*e.g.*, Sætre et al., 1997; Noor, 2000; Naisbit et al., 2001; Yukilevich, 2012; Dyer et al., 2014), decreased species divergence (*e.g.*, Ritchie et al., 1989; Shurtliff et al., 2013; Yang et al., 2016), local extinction of one species due to reproductive exclusion (Hochkirch et al., 2007, Groning and Hochkirch, 2008), or no change in either direction (Abbott et al., 2013). Because *E. anna* and *E. carunculatum* produce reproductively disadvantaged hybrids (Barnard et al., 2017), selection is expected to favor increased premating isolation when the species are sympatric. Within each species, we predicted that female sensilla traits in sympatric populations would diverge from those of allopatric populations indicative of a shift in female preferences to avoid mating with heterospecifics. Contrary to this prediction, sympatric and allopatric *E. carunculatum* populations were not significantly different in mean sensilla trait values (Fig. 2) or sensilla density distributions (Fig. 4E).

Although we observed a trend toward more sensilla in sympatric *E. anna* populations relative to allopatric populations (Figs. 2A, 3A), it is difficult to conduct a robust comparison for this species because *E. anna’s* entire geographic range overlaps with *E. carunculatum*’s range and *E. anna* are often relatively rare (Acorn, 2004; A. Barnard, personal obs.). It was therefore difficult to collect sufficient *E. anna* samples from populations that do not co-occur with *E. carunculatum.* We might, however, expect a stronger pattern of RCD in sympatric *E. anna* females because *E. carunculatum* males can take them in tandem relatively easily, whereas *E. anna* males are typically unsuccessful at taking *E. carunculatum* females in tandem (Barnard et al., 2017). This means that *E. anna* females may have more opportunities for mating mistakes than *E. carunculatum* females, which can result in stronger asymmetric RCD (Lemmon, 2009; Pfennig and Pfennig, 2009).

There are at least three potential explanations for the absence of RCD in the form of significant differences in the sensilla traits we measured between sympatric and allopatric populations of *E. carunculatum.* First, species-specific sensilla distributions may be sufficiently different to allow females to recognize when they are taken in tandem by heterospecific or conspecific males. If this is true, small degrees of variation within the overall species pattern among females might not affect females’ species-recognition abilities. Indeed, a recent study found that intraspecific variation in male cercus morphology appears too minor for *Enallagma* females to show strong discrimination among conspecific males that grasp them (Siepielski et al., 2018). Although RCD is most easily facilitated when the trait under selection already differs between species (Pfennig and Pfennig, 2009), these sensilla traits may have already diverged sufficiently enough to preclude strong selection on further divergence.

Second, it is possible that the external sensilla phenotypes we measured are not representative of proximate female sensory traits, and the variation that directs mating decisions occurs within the female nervous system. For example, individual sensilla might differ in response rate or ability to distinguish different levels of pressure applied by the cerci and grasping pressure might differ between males of each species. The direction of mechanosensor deflection is also important for stimulus detection (Keil, 1997), and different species’ cercus morphologies may contact sensilla from different angles. Female mate preferences may also be influenced by the relative frequencies with which females encounter heterospecific and conspecific males and female sexual experience (*e.g.*, Svensson et al., 2014).

Finally, although we did not detect a statistically significant difference between group means, the small differences we observed may still have biological relevance. If gaining just one additional mechanosensor can (at least) double a female’s tactile discriminatory power (Gaffin and Brayfield, 2017), then females in a population with a seemingly minor upward shift in sensilla number could gain a substantial increase in their ability to detect and avoid mating with heterospecifics. Similarly, it is difficult to determine the features of sensilla density distributions that may influence female preference solely by conducting statistical tests between KDEs. Small spatial differences within largely similar patterns may not contribute a signal large enough to be captured in a statistical test, but still reflect salient variation in the way females receive tactile stimuli. This might include three dimensional spatial differences that we were unable to measure here.

These possible explanations highlight the interesting avenues that female damselfly sensilla provide for investigating the mechanisms underlying how females evaluate male tactile signals to make mating decisions. The ability to quantify the number and locations of female mechanoreceptors in a region contacted by male reproductive structures complements our understanding of patterns of variation in male morphologies (McPeek et al., 2008; McPeek et al., 2009; McPeek et al., 2011; Barnard et al., 2017). Females of both species display substantial intrapopulation variation in sensilla traits (Fig. 3) and this variation may play a role in sexual selection and female preferences within species. Behavioral studies will be crucial to link mechanoreceptor phenotypes to female mating decisions and clarify how sensilla traits influence both species recognition and sexual selection. For example, do females with more sensilla make fewer mating mistakes than females with fewer sensilla (Lemmon, 2009)? Another outstanding question of this system is how the cerci stimulate individual sensilla during tandem. This might be determined by flash-freezing male-female tandem pairs and using micro-CT scanning to understand how the male and female structures interact, similar to a recent approach used in seed beetles (Dougherty and Simmons, 2017). Once we understand how cerci contact the sensilla, functional tests of sensilla electrophysiology could reveal how individual sensilla respond to stimulation and indicate whether certain sensilla make greater contributions to reproductive decision-making than others.

Female preference can drive sexual selection, promote trait divergence, and cause RI between species (Ritchie, 1996). A longstanding presumption in the literature on genital evolution and speciation has been that female reproductive morphologies are less variant or species-specific than male genitalia (reviewed in Shapiro and Porter, 1989). However, recent studies of variation in female reproductive structures suggest that variation does exist among individuals and species (Ah-King et al., 2014), and our data highlight the importance of looking beyond the easily-quantified external morphologies. When male reproductive structure morphologies are obviously divergent, but female morphologies are not, females may possess important variation at neurophysiological levels that affects how they evaluate male tactile signals, similar to the way females evaluate signals in other sensory modalities.

## Acknowledgements

We are grateful to P. Larson for SEM imaging help. We thank O. Fincke, D. Paulson, and B. Mauffray for generously donating damselfly specimens, and D. Gaffin, O. Fincke, R. Knapp, G. Wellborn, and S. Westrop for helpful discussion during the course of this work. This work was supported by funds from the University of Oklahoma to JPM and AAB. The authors declare that there is no conflict of interest regarding the publication of this article.

## Figure legends

**Figure S1.**
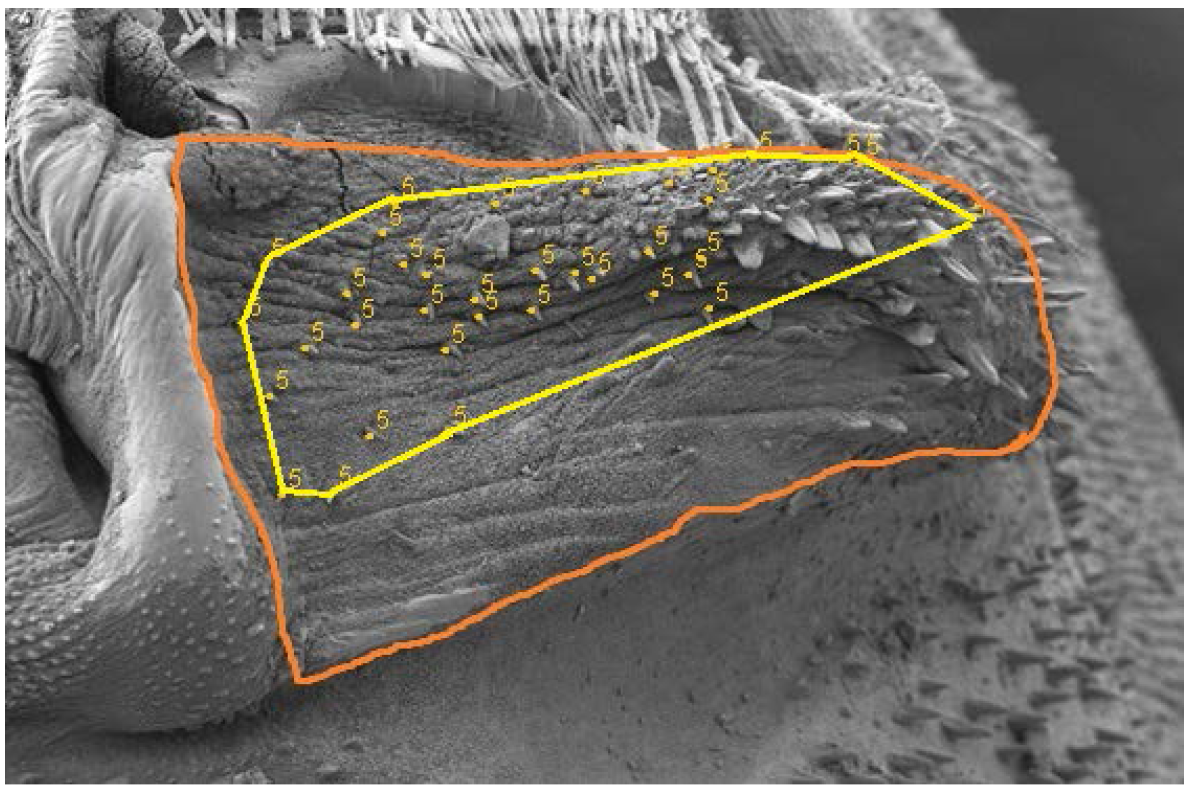
Method to obtain (*x*, *y*) coordinates of mesostigmal plate outline and individual sensilla from scanning electron microscope images. The orange line shows the outline that represents the boundaries of the mesostigmal plate. Yellow dots indicate individual sensilla. The yellow line around the sensilla shows the polygon generated by connecting the outermost sensilla.

**Table S1.**
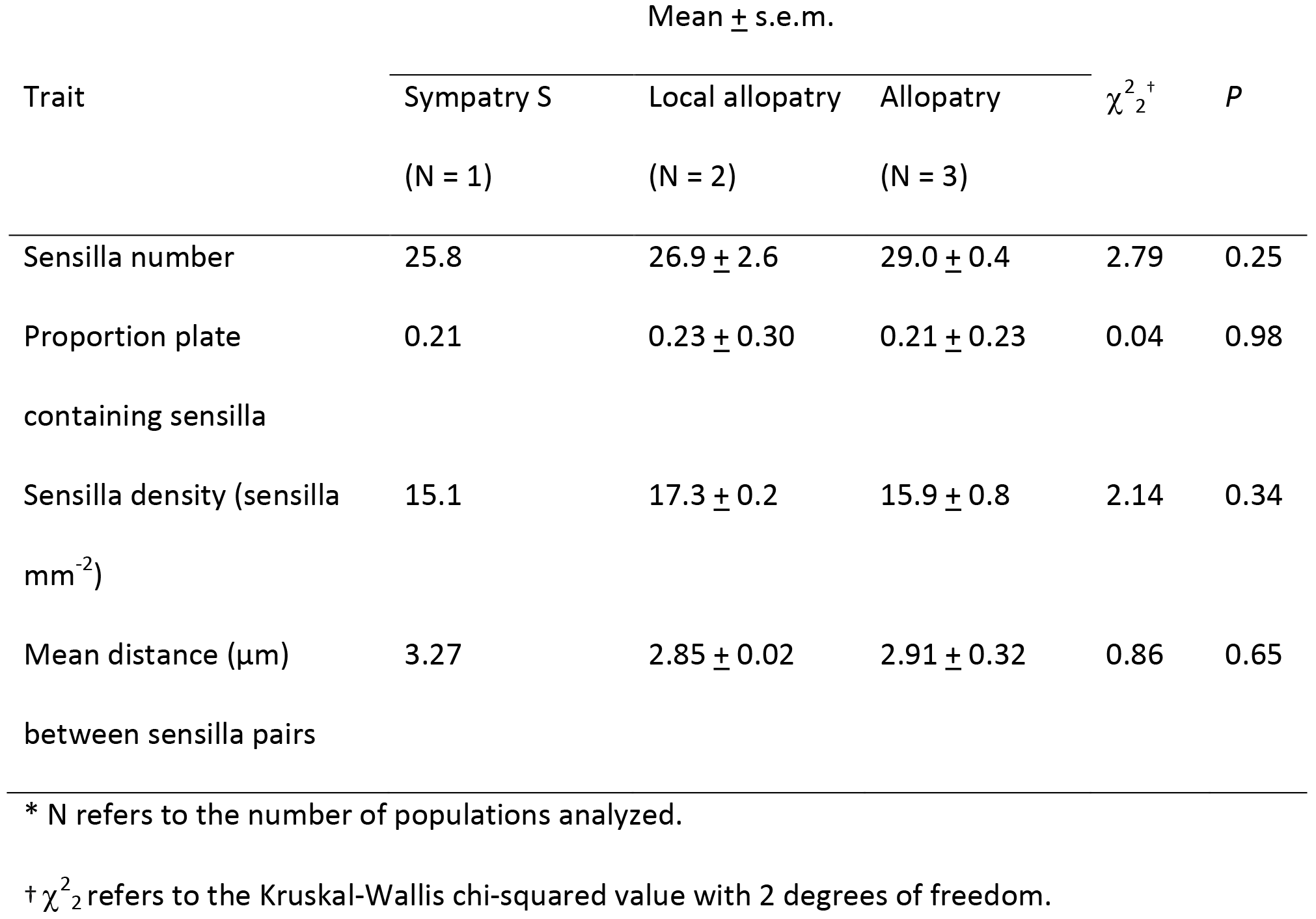
Statistical comparison of sensilla traits among sympatric, locally allopatric, and fully allopatric *E. carunculatum* populations.

